# Relationships among evolutionary distance, the variance–covariance matrix, multidimensional scaling, and principal component analysis

**DOI:** 10.1101/2022.03.02.482744

**Authors:** Kazuharu Misawa

## Abstract

Principal component analyses (PCAs) are often used to visualize patterns of genetic variation in human populations. Previous studies showed a close correspondence between genetic and geographic distances. In such PCAs, the principal components are eigenvectors of the data’s variance-covariance matrix, which is obtained by a genetic relationship matrix (GRM). However, it is difficult to apply GRM to multiallelic sites. In this paper, I showed that a PCA from GRM is equivalent to multidimensional scaling (MDS) from nucleotide differences. Therefore, a PCA can be conducted using nucleotide differences. The new method provided in this study provides a straightforward method to predict the effects of different demographic processes on genetic diversity.

## 1. Introduction

Understanding the structure of human populations is important for human genetic studies. Principal components analysis (PCA) was first applied to genetic data by Cavalli-Sforza and colleagues ^1^. PCAs were widely used to reveal the present patterns of human genetic geography and revealed several hidden patterns of genetic variation in human populations ^1–7^. Previous studies, such as ^4,7,8^, showed that geographical maps naturally reflect a two-dimensional summary of genetic variation. For example, Novembre and his colleagues ^4^ showed a close correspondence between genetic and geographic distances in Europe. Moreover, McVean ^9^ showed that the projection of samples onto principal components can be directly inferred by considering the average coalescent times between pairs of haploid genomes.

The principal components are eigenvectors of the data’s variance–covariance matrix, which is obtained by a genetic relationship matrix (GRM). However, it is difficult to apply a GRM to multiallelic sites. The aim of this study was to develop a method to facilitate PCA using nucleotide differences. Therefore, I investigated the relationship between PCA and nucleotide differences.

## 2. Materials and Methods

I first briefly outline the theoretical formulation of PCA based on McVean’s notation ^9^. I assumed that *N* haploid individuals were sequenced. Let us define a new matrix X with row sums equal to 0.

### 2.2. Variance Covariance Matrix and Genetic Relationship Matrix

The principal components are eigenvectors of the data’s covariance matrix. The first principal component can equivalently be defined as a direction that maximizes the variance of the projected data. Let us define a matrix ***M**_s_* at locus *s*. The *ij*-th element of ***M**_s_* is defined using the following equation:

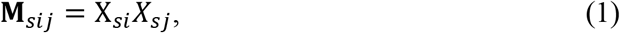

The second step of PCA is to obtain eigenvectors of the matrix **M**. Let us denote the *i*th eigenvalue and eigenvector of the matrix **M** by *λ_i_* and *u_i_*.

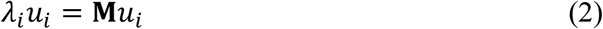

### 2.2. Genetic Relationship Matrix

In the following section, I assumed that all polymorphic sites were biallelic; that is, they have two alleles that are the result of a single mutation. Let *Z_si_* be the allelic state for individual *i* at locus *k*, coding the ancestral allele as 0 and the derived allele as 1. However, the following also applies for any coding, for example the minor allele coded as 1, as mentioned by McVean ^9^. **Z** consists of an *L*×*N* binary matrix where *L* is the number of SNPs.

The vector ***z**_s_* and **1** are defined by:

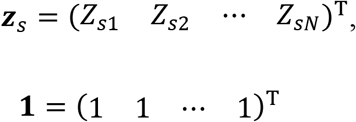

The GRM is obtained as 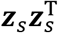 using the genotype valued decomposition^10^. The GRM is used in the software GCTA ^11^ and in PCA ^3^.

The variance–covariance matrix is also obtained by **1** and ***z**_S_*.

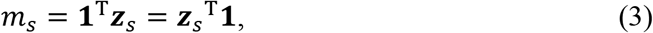

where

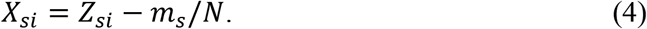

**M**_s_ can be obtained by the following equation:

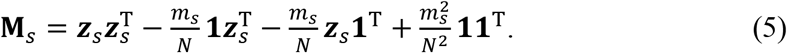

The variance–covariance matrix of the whole genome is obtained as follows:

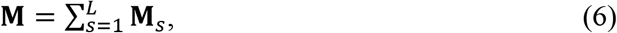

These equations are essentially the same as equations (8)–(10) proposed by McVean ^*9*^.

### 2.3. Evolutionary Distance

I defined a new variable, *d_sij_*, as 1 when the allelic states of *i* and *j* at locus *s* differed; otherwise, *d_sij_* was defined as 0. Because *Z_Si_* and *Z_Sj_* are either 0 or 1, when they both equal 1, then *Z_Si_Z_Sj_* = 1; otherwise, *Z_Si_Z_Sj_* = 0.

D is the number of different sites between the haploid samples *i* and *j*. Let us assume the additivity of evolutionary distances. I obtained the evolutionary distance as follows:

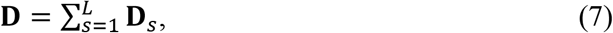

Note the equivalent formula for this defined variable:

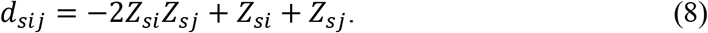

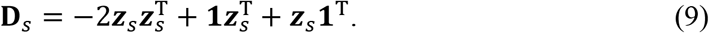

### 2.3. Number of Multiallelic Sites

Number of multiallelic sites were counted from the genetic variation data of 2,504 people determined by 1000G which were downloaded from the International 1000 Genomes Project ^12^.

## 3. Results

### 3.1 Number of Multiallelic Sites

In this paper, I used the double centering of multidimensional scaling (MDS) ^13^, because c the sum of each column is required to be 0 to perform an effective PCA. The following equation was used to perform the MDS ^14^:

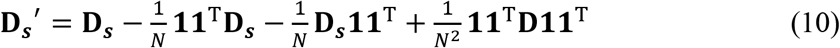

Using equation (5), **D′** is obtained as:

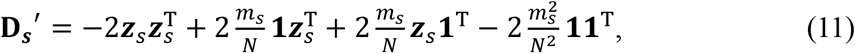

(for details, see Appendix A).

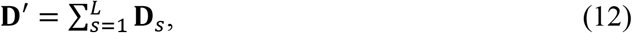

From equations (5), (6), (11), and (12), we obtain

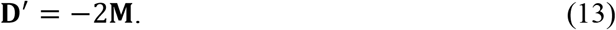

Equation (13) shows a PCA from GRM is equivalent to MDS from nucleotide differences.

### 3.2. Number of Multiallelic Sites

Among genomic sequences of 2,504 people determined by 1000G, 81 million sites were biallelic and 274,620 sites were multiallelic (Table 1).

**Table 1.** Number of SNPs in the 1000G dataset.

| Type | Number of SNPs |
| --- | --- |
| Biallelic | 81,106,364 |
| Triallelic | 272,987 |
| Tetraallelic | 1,633 |
| Multiallelic sum | 274,620 |

## 4. Discussion

I showed that principal components can be directly obtained from the evolutionary distances of haploid genomes.

When GRM among individuals is used for PCA, the GRM is obtained by

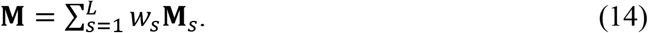

where *w_s_* is weight of locus *s*. Weights are usually included to normalize the data in each column to have the same variance ^2,3^. McVean showed that normalization has little effect upon SNP data ^9^.

A close correspondence between genetic and geographic distances indicates that the geographical distance between two persons is a linear function of the time to their common ancestor. The relationships among migration, geographical isolation, and admixture have also been studied on the basis of a modest number of discrete demes ^7,15,16^. McVean ^9^ showed that the projection of samples onto the principal components can be obtained directly by considering the average coalescent times between pairs of haploid genomes. His result provided a direct route to understanding the influence that various demographic scenarios can have on the relationships between samples identified from PCA.

Moreover, Charlesworth et al ^17^ reviewed some of the theory and provided expected coalescence times, which provided new insight into continuous human movement.

Continuous isolation-by-distance models have been studied to obtain the probability of identity-by-descent ^18–20^ and to study the relationship between gene frequencies and geographic distances ^21,22^ Previous theoretical studies showed that coalescent times vary with deme location ^23,24^ Further studies must be necessary. However, it is worth noting that the evolutionary distance between two randomly chosen genes increases almost linearly with coalescent time, and coalescent time refers to the time to most recent common ancestor.

I assumed polymorphic sites were biallelic. However, in the datasets with large sample sizes, many variant sites were observed to be multiallelic. It is difficult to apply GRM to multiallelic sites, but evolutionary distances can easily be calculated using multiallelic sites. PCA using evolutionary distances will be widely applicable for studying genome variation.

In this study, I assumed additivity of evolutionary distance. However, there are many causes of nonlinear evolutionary distances ^25^. When evolutionary distance is not linear, PCA needs to be conducted using a nonlinear dimension reduction approach ^26^.

## Author Contributions

K. M. confirms sole responsibility for the following: study conception and design, data collection, analysis and interpretation of results, and manuscript preparation.

## Acknowledgments

I thank Prof. Fujio Tajima and Prof. Naomichi Matsumoto for their advice and encouragement. I thank Mallory Eckstut, PhD, from Edanz (https://jp.edanz.com/ac) for editing a draft of this manuscript. This work was supported by JSPS KAKENHI Grant Numbers JP17K08682, JP19K22647, JP20K07316. This work was a part of the branding program as a world-leading research university on intractable immune and allergic diseases supported by the Ministry of Education, Culture, Sports, Science and Technology of Japan.

## Conflicts of Interest

The author declares no competing interests.

## Appendix A

From equation (5), it is easy to obtain equations (A1), (A2), and (A3):

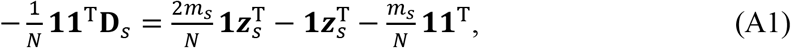

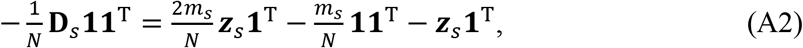

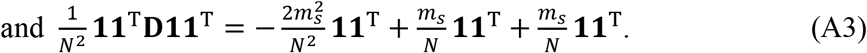

By adding equations (9), (A1), (A2), and (A3), I obtained equation (11).

## Notes

### Competing Interest Statement

The authors have declared no competing interest.

